# Alterations in rhythmic and non-rhythmic resting-state EEG activity and their link to cognition in older age

**DOI:** 10.1101/2021.08.26.457768

**Authors:** Elena Cesnaite, Paul Steinfath, Mina Jamshidi Idaji, Tilman Stephani, Deniz Kumral, Stefan Haufe, Christian Sander, Tilman Hensch, Ulrich Hegerl, Steffi Riedel-Heller, Susanne Röhr, Matthias L. Schroeter, A. Veronica Witte, Arno Villringer, Vadim V. Nikulin

## Abstract

While many structural and biochemical changes in the brain have been previously associated with aging, the findings concerning electrophysiological signatures, reflecting functional properties of neuronal networks, remain rather controversial. To try resolve this issue, we took advantage of a large population study (N=1703) and comprehensively investigated the association of multiple EEG biomarkers (power of alpha and theta oscillations, individual alpha peak frequency (IAF), the slope of 1/f power spectral decay), aging, and aging and cognitive performance. Cognitive performance was captured with three factors representing processing speed, episodic memory, and interference resolution. Our results show that not only did IAF decline with age but it was also associated with interference resolution over multiple cortical areas. To a weaker extent, 1/f slope of the PSD showed age-related reductions, mostly in frontal brain regions. Finally, alpha power was negatively associated with the speed of processing in the right frontal lobe, despite the absence of age-related alterations. Our results thus demonstrate that multiple electrophysiological features, as well as their interplay, should be considered when investigating the association between age, neuronal activity, and cognitive performance.

## 1. INTRODUCTION

Older age is often associated with cognitive decline that is accompanied by structural changes in the brain [1]–[3]. Such changes could also be accompanied by alterations in synchronous firing of pyramidal neural cells involved in the generation of rhythmic oscillatory activity that can be measured with scalp EEG [4]–[7]. Oscillations are typically defined by power, peak frequency, and phase. While all of these parameters have been previously linked to cognition, alterations in power and peak frequency of different oscillatory bands have been also extensively studied in aging [8]– [12]. However, a number of contradictory findings (presented below) suggest that the association between these parameters, age, and cognitive decline remain unclear.

Previous work has shown that theta and alpha oscillations (approx. 4 - 7Hz and 8 - 12Hz, respectively) play an important role for cognitive functions [7], [11]–[13]. Their involvement in higher cognitive processes has been explained through the top-down control over processed information. An interplay between the oscillatory activity in alpha and theta frequency bands enables suppression of task irrelevant information that helps direct attention towards task relevant stimuli [10], [12], [14], [15]. While a decrease in alpha power has been suggested to reflect release from inhibition, theta power increase might facilitate encoding of new information [11], [16], [17]. Functional significance of the power and peak frequency of theta and alpha oscillations is supported by their link to structural and biochemical alterations in the brain, which is most prominent in older age (above 60 years; [18]–[21]. While most studies consistently show age-related slowing of individual alpha peak frequency (IAF; [9], [22], [23], previous findings are inconsistent regarding changes in oscillatory power of theta and alpha oscillations. While alpha power has been suggested to either decline with age [24] or show no age-related alterations [25], theta power has been shown both to decline [26] and increase [11], [21], [22], [27] with age.

On the one hand, these inconsistencies might be due to power estimation in canonical, rigidly pre-defined frequency bands: if a possible center frequency shift is not accounted for (i.e., in a case of spectral slowing), power estimation of two signals in different frequency ranges might be confounded. For instance, it has been suggested that slowing of an IAF might interfere with the conventional theta frequency range and thus result in its estimated power increase due to the interference of an alpha peak [28]. On the other hand, slow-wave power estimation is complicated not only due to center frequency shift but also due to the amplitude mixing between the oscillatory and non-oscillatory (i.e., non-rhythmic) activities [29]–[31]. Non-rhythmic activity results from asynchronous spiking and postsynaptic potentials of neural populations and it can be estimated with the 1/f slope of the power spectral density (PSD, [30], [32]). Traditionally considered as noise, it has only recently gained interest as its functional significance becomes more recognized [32]– [35]. The aperiodic component of the PSD has been previously linked to cortical states reflecting the ratio between excitatory and inhibitory inputs, as determined by glutamatergic and GABAergic connections respectively [30], [35], [36]. An increase in excitatory connections was associated with a power increase in higher frequency ranges and, therefore, a flatter PSD slope, as compared to an increased number of inhibitory connections, which resulted in a steeper PSD slope. This relationship has been further supported by pharmacological intervention studies of altered states of consciousness [37], [38]. Age-related changes in 1/f slope have been addressed in a few previous studies showing that the slope decreases as we age (i.e., flattens), suggesting an increase in excitability and neural noise [31], [33]. Importantly, when controlling for 1/f decay of PSD (decomposing oscillatory and non-oscillatory estimates of PSD), no age-related alterations were shown in slow wave (< 12 Hz) power [25]. Furthermore, few studies have investigated the association between non-oscillatory activity estimated with 1/f slope and cognition, being linked to cognitive speed [32], lexical prediction [33], and visual working memory [31].

In the current study, we aimed to disentangle the complex relationship between oscillatory and non-oscillatory EEG parameters, age, and cognition in a large elderly sample from a population-based study (LIFE-Adult; [39]). The hope was to identify biomarkers that could identify preserved cognitive function in older age. As periodic (i.e., alpha power, IAF) and aperiodic (i.e., 1/f slope) components of PSD have been shown to relate to cortical excitability and several cognitive functions, we aimed to investigate their individual contributions in determining the relationship between the proposed measures. We hypothesized that no age-related alterations in theta and alpha power would be observed when carefully adjusting for methodological confounds, since alterations previously reported in the literature could have been a result of spectral slowing or amplitude mixing. Moreover, based on previous literature, we have also hypothesized a decrease of IAF and 1/f slope with increasing age. Regarding the functional significance of such alterations, we expected that higher IAF and higher 1/f slope values would relate to better cognitive performance, based on previous findings [31], [40].

## 2. RESULTS

### 2.1. Descriptive information

Demographic information and sample characteristics can be found in *Table 1* separately for men and women.

**Table 1.** Sample descriptive information separately for men and women. Abb.: IAF – individual alpha peak frequency, SD – standard deviation

|  | *Men (n=823)* |  |  |  | *Women (n=880)* |  |  |  |
| --- | --- | --- | --- | --- | --- | --- | --- | --- |
|  | Min | Max | Mean | SD | Min | Max | Mean | SD |
| *Age* | 60 | 79.6 | 70.3 | 4.6 | 60 | 80 | 69.9 | 4.8 |
| *Education (years)* | 10 | >17 | 14 | - | <10 | >17 | 12 | - |
| *Theta power($\mu V^2$)* | 0.12 | 9.19 | 1.18 | 1.11 | 0.16 | 25.67 | 1.61 | 1.71 |
| *Alpha power($\mu V^2$)* | 0.47 | 109.44 | 17.84 | 15.65 | 0.56 | 207.87 | 22.12 | 23.58 |
| *IAF (Hz)* | 7.33 | 12.06 | 9.29 | 0.86 | 7.63 | 12.59 | 9.61 | 0.88 |
| *1/f slope* | 0.55 | 1.82 | 1.18 | 0.19 | 0.16 | 1.73 | 1.15 | 0.21 |

*Figure 2* shows the PSD of an exemplary single channel and an aperiodic fit (derived from the FOOOF algorithm) as well as the scalp topographies of the grand average of the rsEEG parameters across subjects. The power of alpha band oscillations shows a typical occipital distribution of higher values and is comparable to the topography of theta power, which might be indicative of an absence of oscillatory theta activity. The topography of 1/f slope shows that it has higher values around the midline.

**Figure 1.**
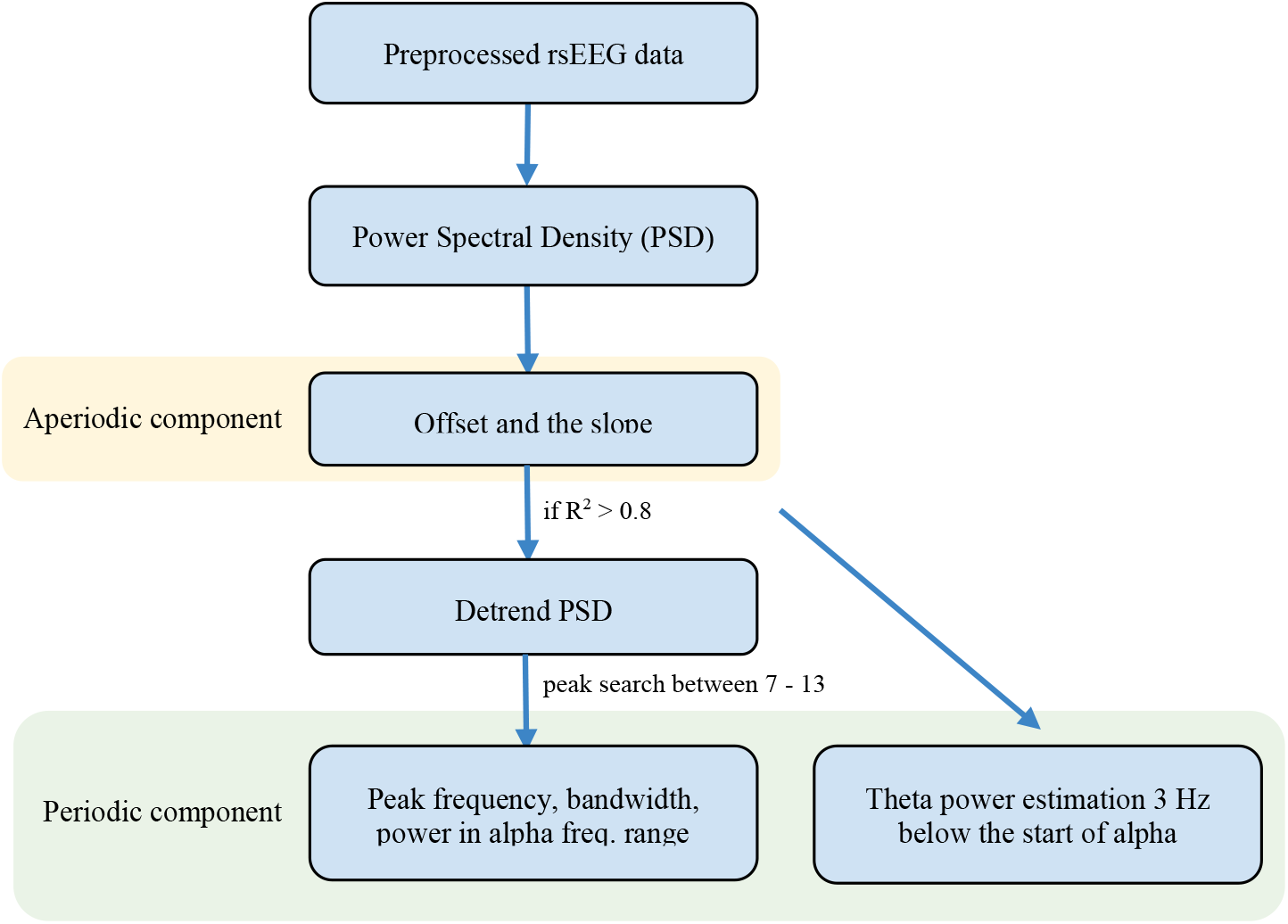
Schematic depiction of the rsEEG analysis pipeline decomposing power spectral density into aperiodic and periodic components. We observed a consistent presence of the oscillatory peak only in the alpha frequency range. Theta power was therefore estimated using the original PSD in the 3Hz range prior to the starting frequency of the alpha peak.

**Figure 2.**
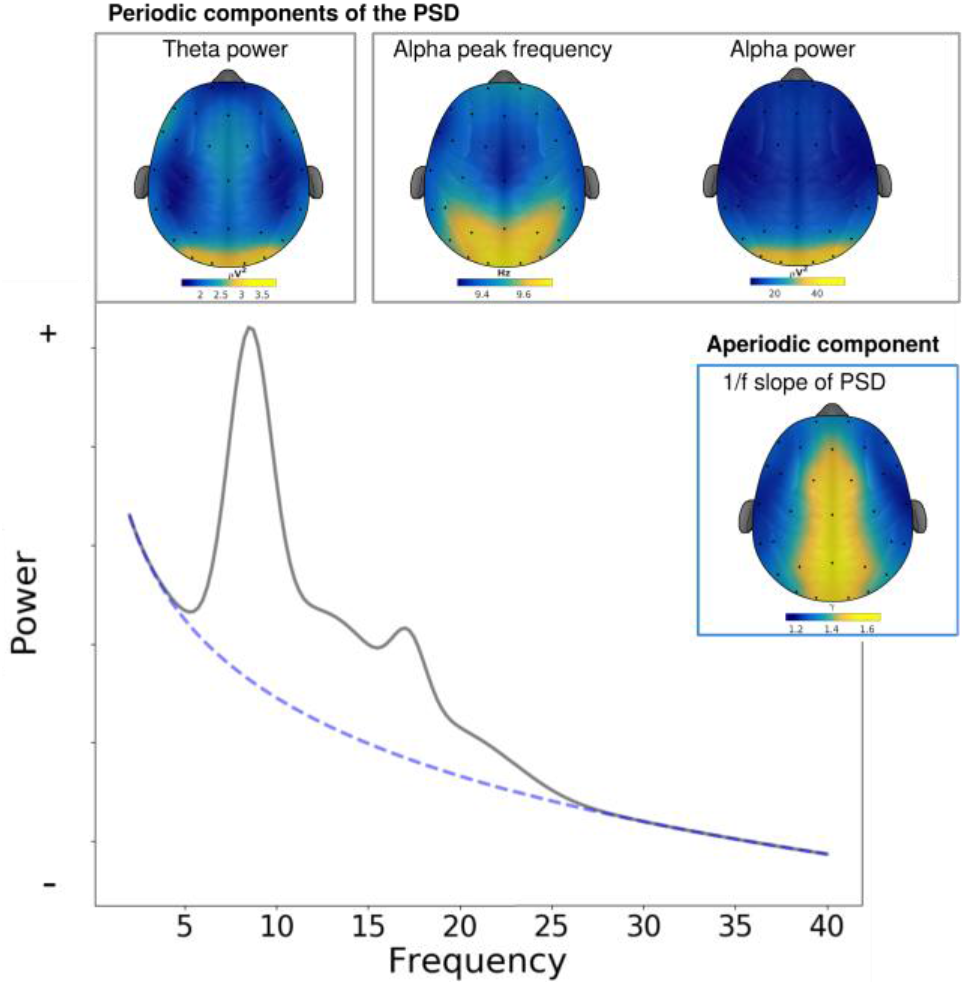
Grand average topographies of resting-state EEG parameters and an exemplary power spectral density (PSD) of a single EEG channel. Periodic parameters of theta and alpha oscillations were estimated in individually defined frequency ranges while the aperiodic component was estimated between 2 and 40 Hz and is indicated here by 1/f slope of the PSD (dashed line).

Theta and alpha power, IAF, and 1/f slope of PSD showed strong correlations among each other with a widespread effect over the whole cortex (p-values were based on Pearson correlation, clusters of electrodes where a significant relationship was found were corrected for multiple comparisons using cluster statistics, *Figure 3*).

**Figure 3.**
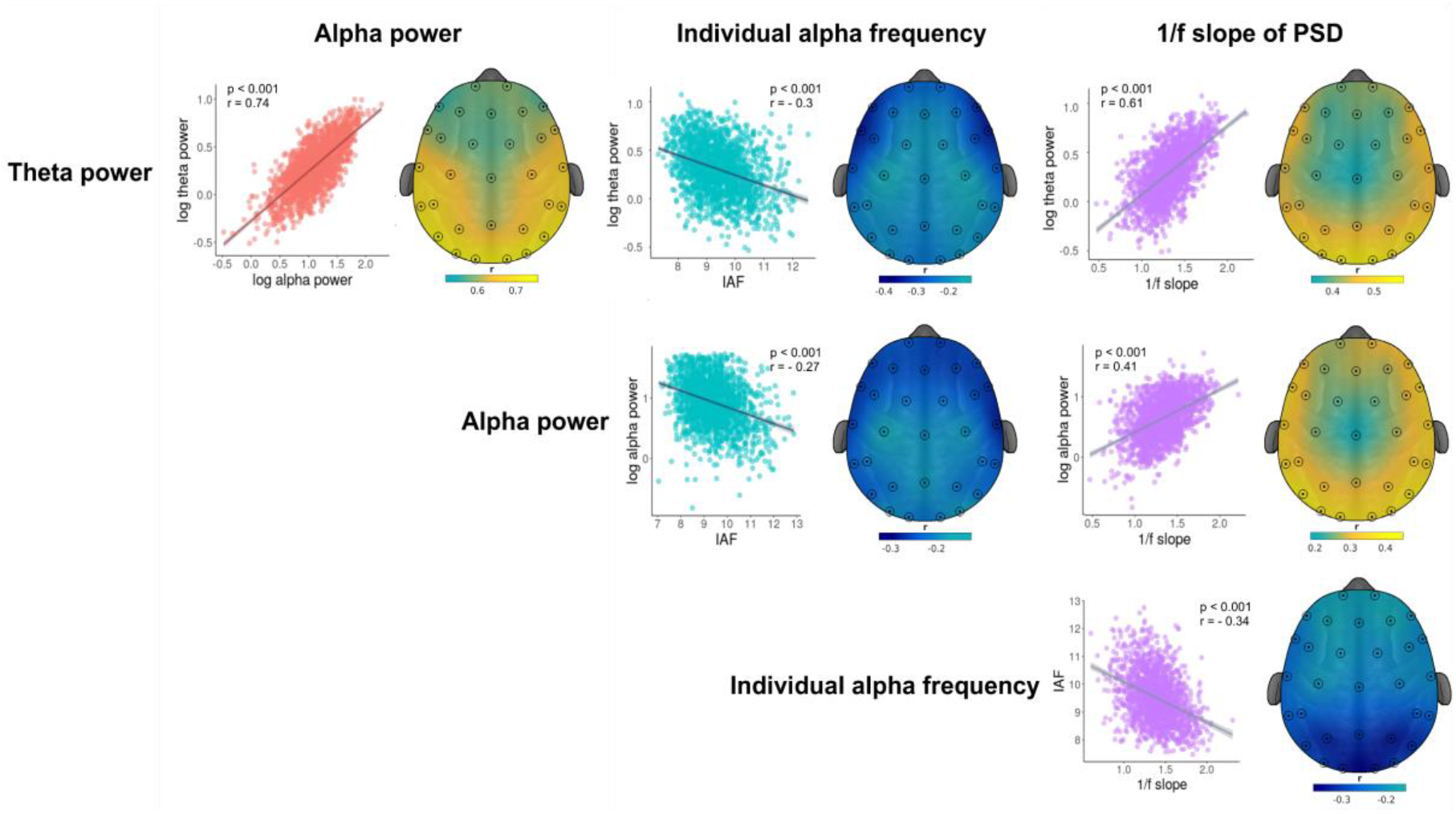
Correlation between periodic and aperiodic components of the PSD. Scatter plots show linear relationships between the parameters included in the study averaged across electrodes. Topographical distributions of these relationships reveal clusters of electrodes where the significant relationship between the two measures was found. Sensors that formed a significant cluster are marked with circled black dots.

### 2.2. Resting state EEG spectral changes associated with age

We used a mass-bivariate approach with cluster-based permutation tests to assess whether age-related changes occur in the rsEEG parameters of interest at the sensor space. Topographies representing significant clusters of age-related alterations in theta and alpha power, IAF, and 1/f slope can be seen in *Figure 4*. A widespread negative relationship between IAF and age (p_*cluster*_ < 0.001, *Figure 4, panel (C)*) was present at all electrodes. This result was replicated at the source level, where this relationship was significant in all 10 ROIs with the left temporal lobe showing the strongest effect (p < 0.001, r = -0.22; *Figure 4, panel (C)*). Another cluster of electrodes showing a negative significant relation between 1/f slope and age was detected over the fronto-central electrodes at sensor space (p_*cluster*_ = 0.012, *Figure 4, panel (D)*). Source reconstruction showed that the significant relationship was detected in the right frontal lobe (p = 0.001, r = -0.08; *Figure 4, panel (D)*).

**Figure 4.**
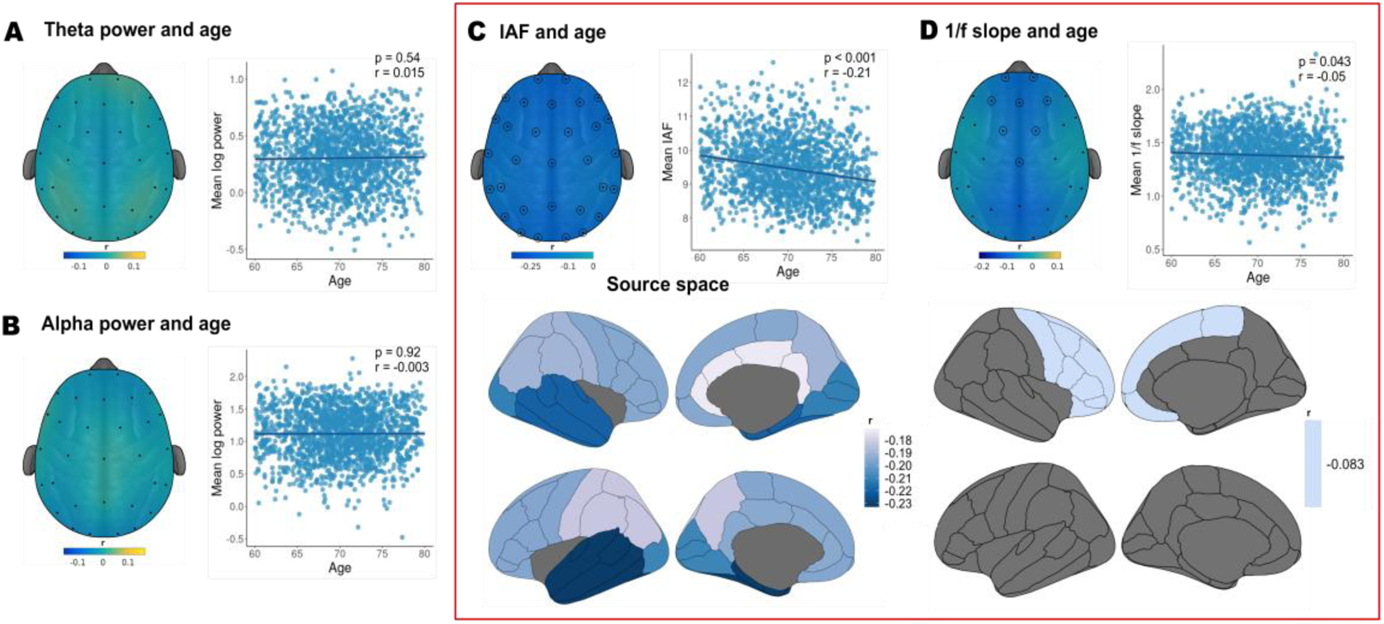
Age-related alterations in resting-state EEG parameters as shown by topographies of correlation coefficients and scatterplots of mean values across EEG sensors. Sensors that formed a significant cluster are marked with black circled dots. No significant relationship was observed between (A) age and theta power, as well as, (B) age and alpha power. (C) Individual alpha peak frequency (IAF) showed a significant negative correlation with age that was prominent over the whole cortical mantle at sensor and source space. (D) 1/f slope of the power spectral density was found to be negatively associated with age: a result that has been detected over the fronto-central channels at sensor space and right frontal lobe at source space. Two significant results are indicated by a rectangle with red borders.

Other rsEEG parameters of interest such as theta power, and alpha power showed no significant age-related alterations, neither at sensor nor at source space. Because sensor space findings matched findings in the source space, source space data were used in the following analyses.

### 2.3. Cognitive performance

#### 2.3.1. Factor analysis

We used exploratory factor analysis (EFA) to extract latent factors underlying the cognitive tests. Based on the factor loadings (higher than 0.4), we interpreted the three factors as representing *speed of processing, episodic memory*, and *interference resolution* (see *Figure 5, panel (A)*). The first factor, interpreted as the speed of processing, positively loaded on the reaction times from the congruent and incongruent trials of the Stroop task. The second factor, associated with episodic memory, positively loaded on the WMS Logical Memory Scale and the number of details recalled before and after the break. Finally, the third factor was interpreted as reflecting interference resolution since it loaded positively on the accuracy of incongruent trials of the Stroop task and negatively on the inverted reaction times of the same task. The interpretation of the third factor as interference resolution was based on the increased cognitive demands under the effect of Stroop interference that results in slower reaction times but higher accuracy. The three identified factors (processing speed, episodic memory, and interference resolution) correspond to the main cognitive domains of attention, memory, and executive functions, respectively.

**Figure 5.**
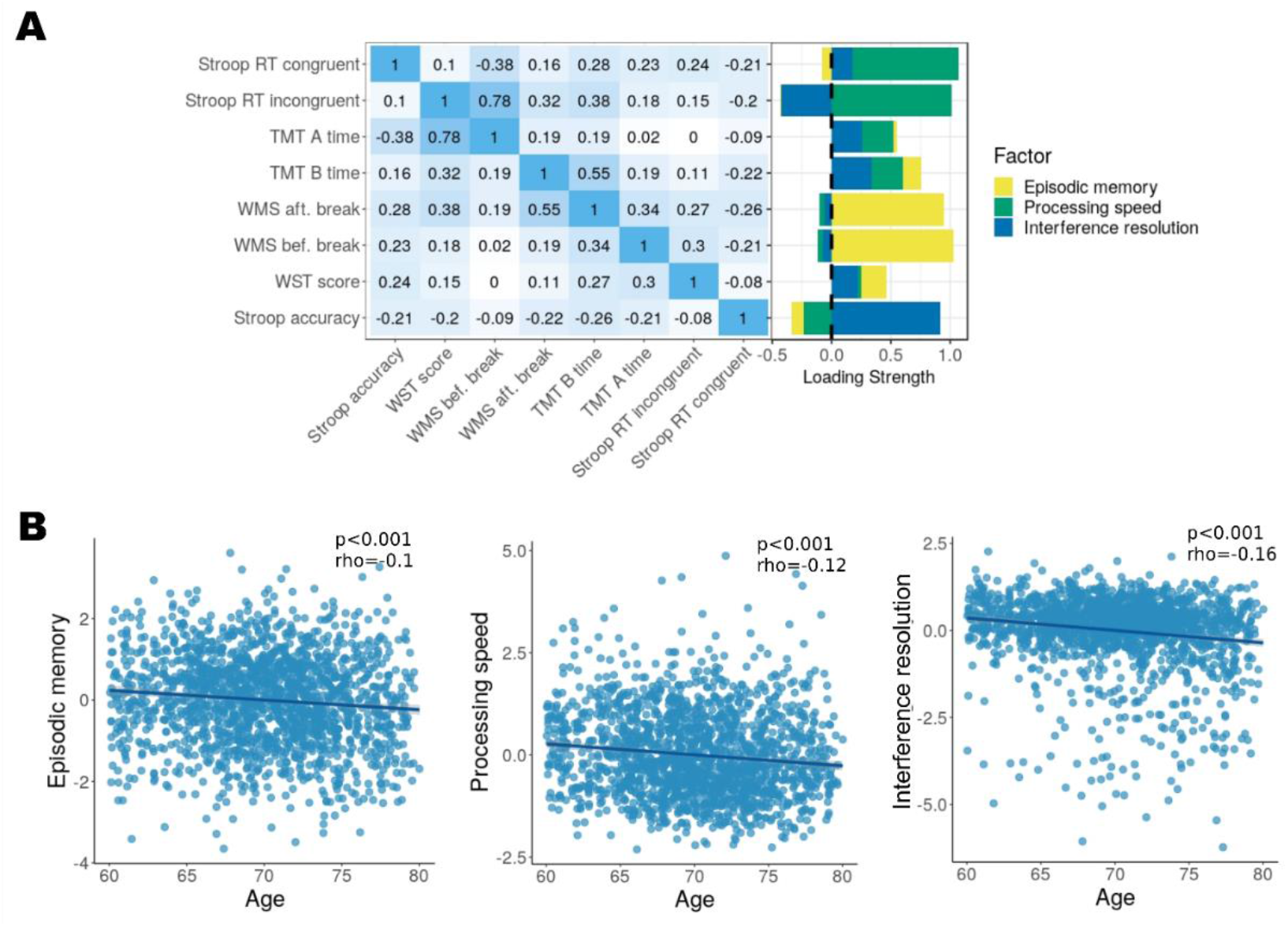
Three latent factors underlined the cognition battery and showed significant associations with age. (A) The correlation matrix shows that most of the subscales were moderately correlated with each other. Three latent factors were derived from the factor analysis: processing speed, episodic memory, and working memory. The loading strength, to the right, represents the contribution of the particular scale to the factor. (B) All of the latent factors showed a decrease with age.

#### 2.3.2. Relationship between age and cognitive performance

All three factors showed a significant decrease with age (speed of processing: p < 0.001, rho = -0.12, episodic memory: p < 0.001, rho = -0.1, interference resolution: p < 0.001, rho = -0.16; *Figure 5, panel (B)*) and were used for the following multiple linear regression (MLR) models.

### 2.4. Relationship between resting-state EEG parameters and cognitive performance

We used MLR to estimate which of the rsEEG parameters were associated with cognitive performance. Models for each cognitive factor derived from EFA were used separately for each ROI. For simplicity, we report here only the results from regions that were statistically significant, all outputs from the MLR models can be found in Supplementary material *Tables 1-10*.

Models for the first factor representing the speed of processing revealed a significant negative relationship with alpha power (p < 0.01, β = -0.08; *Figure 6*) in the right frontal region (*model statistics*: adj. R^2^: 0.017; F(1557) = 3.41, p < 0.001). Age was also negatively associated with the factor in this model (p < 0.001, β = -0.11). Other independent variables as well as interaction terms between rsEEG parameters and age were not significant.

**Figure 6.**
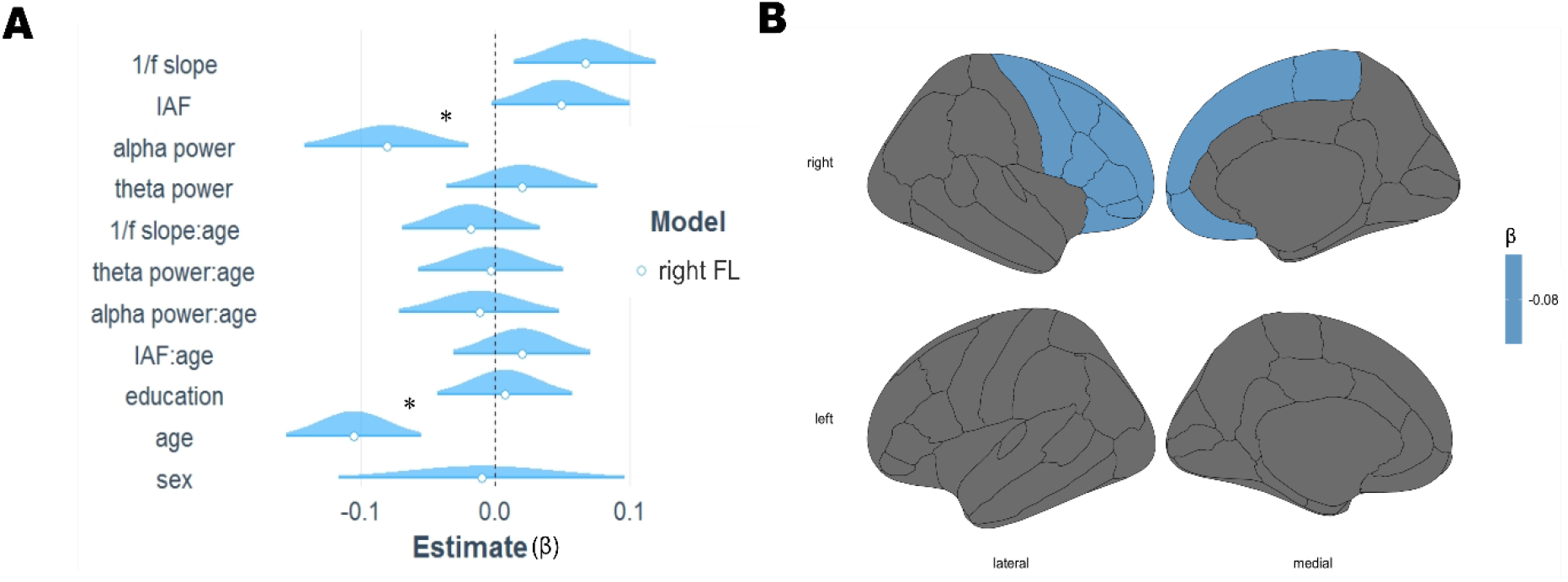
A negative relationship between alpha power in the right frontal lobe and speed of processing. A multiple linear regression model of this region revealed that reduced alpha power was related to faster speed of processing when all other EEG parameters, as well as age, sex, and education, were kept constant. (A) The figure shows estimates of standardized predictors together with a distribution of their confidence intervals. Interaction terms are indicated with a colon and significant effects are marked with an asterisk. (B) The significant region plotted at the source space indicates the relationship between alpha power and processing speed on the cortical mantle.

Models predicting the second factor, representing episodic memory were significant (*model statistics*: adj. R^2^ ranged from 0.061 to 0.068; F(1557) ranged from 10.35 to 11.36, p < 0.001). However, they showed no association with the rsEEG measures.

Models predicting the third factor, representing interference resolution (*model statistics*: adj. R^2^ ranged from 0.03 to 0.04; F(1557) ranged from 5.1 to 7.1, p < 0.001; *Figure 7*) showed a positive association with IAF in six regions, thus indicating that higher IAF was associated with better interference resolution. This relationship was present in the following regions: right frontal (p < 0.01, β = 0.07), right (p < 0.01, β = 0.08) and left (p < 0.01, β = 0.07) parietal, right (p < 0.001, β = 0.09) and left (p < 0.01, β = 0.08) temporal, and right cingulate cortex (p < 0.01, β = 0.07). Moreover, age was negatively related (p < 0.001, βs ranged from -0.14 to -0.15) and education (measured in years) was positively related (p < 0.01, βs ranged from 0.07 to 0.08) to this factor. No other parameters nor interaction terms were significant.

**Figure 7.**
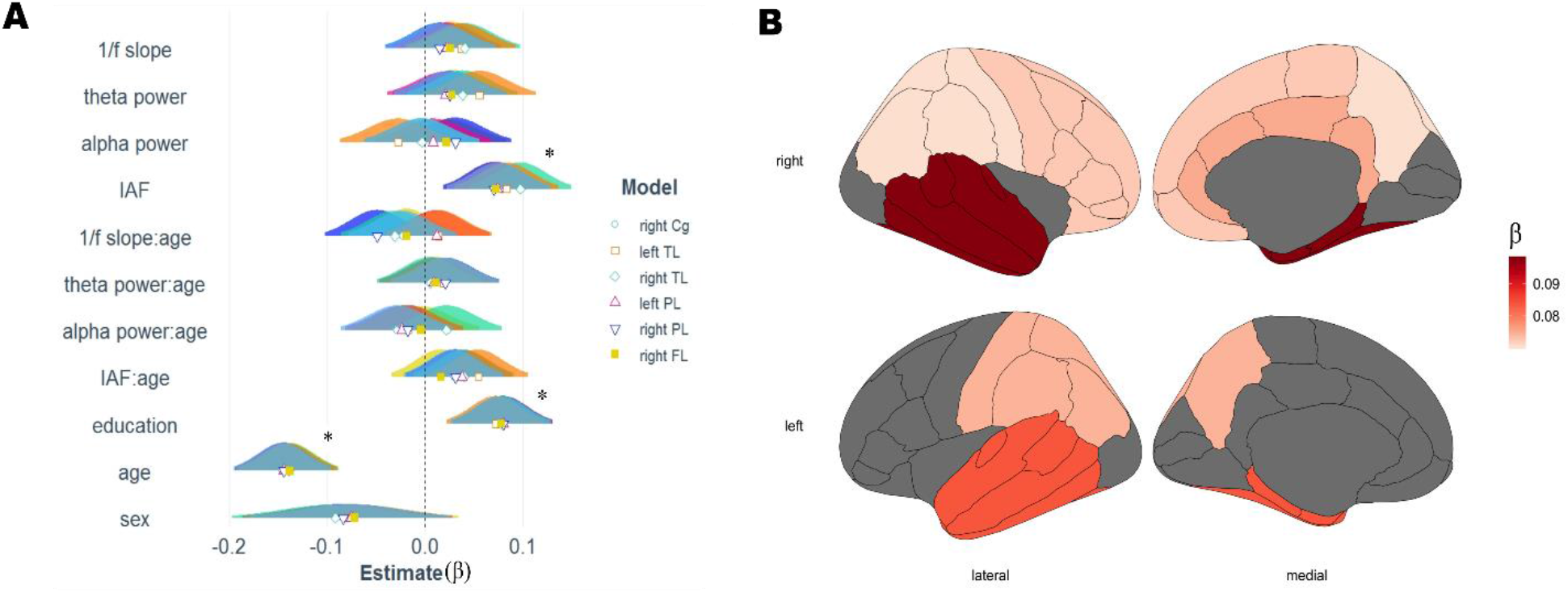
Relationship between individual alpha peak frequency and interference resolution. Multiple linear regression models for each region of interest revealed that IAF was related to the factor representing interference resolution in 6 major regions. (A) The figure shows estimates of standardized predictors together with a distribution of their confidence intervals in different regions indicated by colors. Interaction terms are indicated with a semi-column and significant effects are marked with an asterisk. (B) A positive association between the IAF and interference resolution seen in panel (A) plotted at source space.

## 3. DISCUSSION

This study investigated how rhythmic (*periodic*: theta power, alpha power, IAF) and non-rhythmic (*aperiodic*: 1/f slope) rsEEG activity relates to aging and cognition in large cohort of healthy elderly participants. There were four main findings. (i) IAF decreased with age. an effect that was robustly observed across the whole cortex but was strongest in the left temporal lobe. (ii) Age-related alterations were observed in 1/f slope of PSD suggesting flattening of the slope in the right frontal lobe. (iii) No significant age-related alterations were seen in slow wave power (alpha and theta frequency bands), which is in contrast to several previous reports [21], [22], [24], [26]. (iv) Relating individual contributions of rsEEG parameters to cognitive performance, alpha power in the right frontal lobe was negatively associated with processing speed while higher IAF in multiple cortical areas contributed to better interference resolution.

### Age-related alterations of resting-state EEG parameters

In the current study we show a prominent IAF decrease with age that is consistent with previous reports on age-related spectral slowing [9], [22], [23]. Peak frequency slowing might reflect changes on the level of neurotransmission as well as a decrease in axonal conduction velocity [41], [42]. This might, in turn, result in a prolonged time delay within an intra-cortical circuitry and, therefore, slower IAF [43]. Moreover, lower IAF has been previously considered as a marker of neurodegenerative diseases, such as mild cognitive impairment (MCI; [44]) or vascular dementia [45]. Although participants of the current study did not suffer from MCI or any other neurological conditions, slowing of the alpha peak might nevertheless be indicative of neuronal processes that underlie early subclinical stages of neurodegenerative conditions whose prevalence increase with age. This finding is juxtaposed with the observation that higher IAF might contribute to better cognition, and specifically, interference resolution (see below).

Alterations in axonal connections between neurons can not only affect IAF [46], [47], but can also impact compensatory increases of neural firing rates, resulting in increased power in higher frequency ranges [41] and, therefore, a flatter 1/f slope. In line with these findings, our results show age-related alterations in 1/f slope of the PSD, suggesting an increase in cortical excitability in older age [35]. Importantly, our findings suggesting a link between 1/f slope and age are consistent with previous reports [31], [33] despite differences in analyzed age-ranges. While previous studies compared younger (20-30 years old) and older (60-70 years old) participants, we have observed the same effect within a narrow age range between 60 and 80 years. Observations of 1/f slope alterations have previously been reported in fronto-temporal [31] as well as fronto-central regions [33]. Despite a fronto-central cluster at sensor space in our study, the analysis of source space data showed that the 1/f slope decrease was most prominent over the right frontal lobe. Age-related alterations in the frontal lobe have previously been linked to functional reorganization of the brain, associated with a posterior-anterior shift and a functional ‘over-recruitment’ of frontal brain regions [48], [49] as well as with the neural noise hypothesis [47], [50], which might as well be reflected in the flattening of 1/f slope.

Despite multiple reports of age-related alterations of power in theta and alpha frequency ranges across the lifespan [24], [26], but also within an older age range alone [51], we did not observe significant changes of these parameters. An absence of changes in theta and alpha power might be due to few methodological considerations: Individually adjusted frequency ranges of interest based on the center frequency, the dissociation between periodic and aperiodic components of PSD, and the estimation of unique contributions of these parameters. In line with our findings, Caplan et al [25] did not observe any age-related alterations in theta and alpha band power. While the authors reported a detectable rhythmic activity in the alpha frequency range, it did not alter with age when controlling for non-rhythmic activity (i.e., 1/f slope). This finding suggests that previous reports, showing changes of power in lower frequency ranges with age, might have been due to amplitude mixing between periodic and aperiodic components of the PSD. Another possible explanation for the absence of a significant relationship between the two measures might be structural alterations, such as white-matter lesions, that could mask the aging effect on the oscillatory power [18], [52].

### The link between resting-state EEG parameters and cognition

With the second research question, we aimed to explore the link between rsEEG parameters and cognition in old age. We found that reduced alpha power in the right frontal lobe – when measured at rest – was differentially associated with higher processing speed. Task-related power reduction in the alpha frequency band has been previously linked to an increase in excitability [16], [17], [53]. When measured at rest, alpha power has been suggested to reflect properties of an attentional filter, which might relate to the ability to inhibit task-irrelevant information when task demands are met [54], [55]. In line with these previous studies, we suggest that reduced alpha power over the right frontal lobe might represent increased excitability of a network that enables top-down control. Increased excitability in this region might, therefore, result in impulsive and fast reactions to stimuli at the expense of accuracy (represented by the processing speed) and might serve as yet another support for functional ‘over-recruitment’ of frontal brain regions in older age [48], [49]. Moreover, consistent with our finding, previous reports have also suggested weakened inhibitory activity in older age [42], [56]. While other parameters used in the current study (e.g. IAF and 1/f slope of the PSD) have also been associated with excitability, we have controlled for their possible effects on the respective cognitive function.

It has been shown that not only alpha power but also instantaneous IAF is related to excitation/inhibition balance and information processing, particularly in the visual domain [57]– [59]. The link between IAF and cognition is often estimated in pre-stimulus windows during a task and less so during the resting state. Regarding resting-state activity, Surwillo [60] compared young and old participants and showed a link between the slowing of IAF, increased reaction times, and age. However, we believe that the relationship between age-related alterations of IAF and cognition is more complex and is not solely manifested in the slowing of the reaction times, but also accuracy. While we found a prominent IAF decrease with increasing age, we also observed that IAF was related to working memory demands. Higher IAF over bilateral cingulate cortex, left and right parietal- and temporal-lobes, as well as the right frontal lobe was associated with better interference resolution, and specifically with tasks relying on Stroop interference effects. It has been previously suggested that higher IAF relates to a finer temporal sampling of visual information [59], also shown in a cross-modal domain [58]. The authors have suggested a mechanistic explanation that is based on the number of alpha cycles within which the temporally close stimuli fall – in this way higher IAF might facilitate segregation (in contrast to integration) of discrete perception [59]. Therefore, our findings indicate that while higher IAF corresponds to more oscillatory cycles within the given time window of stimulus presentation, as compared to slower IAF, it might as well facilitate segregation of information resulting from two interfering domains – the color and semantics of the word in the context of a Stroop task.

### Limitations of the study

A few limitations of the current study should be addressed. Longitudinal study design as well as a bigger age range might be important to detect age-related alterations of power in alpha and theta frequency ranges. Moreover, we relied on 10 ROIs based on a coarse parcellation of the cortex at source space due to the small number of electrodes (N = 31). Finally, a finer parcellation would have allowed a better spatial precision of our observations.

### Conclusion

Taken together, our results indicate a negative relationship between IAF and age – a widespread effect that was present across the entire cortical mantle. Further, a positive link between IAF and interference resolution was observed, suggesting that higher IAF may facilitate the segregation of interfering information in elderly age. We also found age-related alterations in 1/f slope of the PSD in the right frontal lobe, consistent with previous studies. Importantly, this finding was replicated in a narrow age range between 60 and 80 years showing a continuous decline in 1/f slope that is in line with the neural noise hypothesis of older age [47], [50].

However, despite previous reports of age-related alterations of theta and alpha power, we did not observe such a relationship in a large sample of typically aging older participants. This could be due to a relatively narrow age range of our participants where the changes in power of oscillations are less pronounced. Yet, our findings are also in line with recent research [25] indicating the importance of dissociating periodic and aperiodic activities. Finally, we also showed that reduced resting-state alpha power in the right frontal lobe was linked to speed of processing, possibly through increased excitability that might lead to impulsive responses to stimuli. Our findings, therefore, are in line with previous research [48], [49] suggesting a possible over-recruitment of frontal brain regions in older age.

## 4. METHODS AND MATERIALS

### 4.1. Participants

The data used in the present study is a part of the population-based LIFE-Adult dataset (Leipzig Research Center for Civilization Diseases, Leipzig University; a more detailed description can be found at [39]). The LIFE-Adult dataset consists of ca. 10,000 mostly elderly participants that were randomly selected from the residence registration office. Invitation letters were sent to potential participants and all participants that agreed to take part in the study provided written informed consent, and received monetary compensation. The study was approved by the ethics board of the Medical Faculty of the University of Leipzig.

EEG data was available from 3390 participants. Our inclusion criteria for the study consisted of completion of cognitive tests (described in section 2.4.), right-handedness, no history of brain hemorrhage, concussion, skull fracture, brain surgery or brain tumor, and no use of medication with an effect on the central nervous system. Participants underwent a structured clinical interview for DSM-IV axis I disorders. We also excluded data from participants that had consumed alcohol or more than a single cigarette on the day of the resting-state EEG (rsEEG) recording. In addition, we required a good data quality (i.e., time-series data was not contaminated by segments of noise that in total lasted for more than half of a recording), 5 min of vigilant resting state recordings (further described in section 2.2.). After exclusion of data that did not meet these inclusion criteria, the total sample consisted of 1703 subjects’ (M_*age*_ = 70, SD = 4.7, 880 females; see *Table 1*) datasets.

### 4.2. Resting-state EEG recordings and pre-processing

A 20min eyes-closed rsEEG data was recorded from 31-channel Ag/AgCl scalp electrodes (Brain Products GmbH, Gilching, Germany) in an electrically shielded and soundproof EEG booth. The electrodes were mounted in an elastic cap (easyCAP, Herrsching, Germany) according to the international standard 10 - 20 extended localization system. The signal was amplified with a QuickAmp amplifier (Brain Products GmbH, Gilching, Germany). Additionally, two electrodes recorded vertical (vEOG) and horizontal (hEOG) eye movements above and beneath the right eye. One bipolar electrode attached to the right and left forearm recorded the electrocardiogram (ECG) activity. The electrodes were referenced to the common average reference with AFz being a ground electrode. The electrodes’ impedances were kept below 10kΩ, the sampling rate was 1000Hz, and the data was lowpass filtered at 280Hz. A more detailed description can be found in a paper by Jawinski and colleagues (2017)[61].

EEG data was pre-processed using the MATLAB-based (MathWorks, Inc, Natick, Massachusetts, USA) EEGLAB toolbox (version 14.1.1b) and custom written scripts. First, the data was band-pass filtered between 1 and 45Hz (4th order Butterworth filter applied back and forth) with a notch filter at 50Hz to remove any remaining power line artifacts. The data was then downsampled to 500Hz. We excluded vEOG, hEOG, and ECG channels from the dataset and visually inspected the PSDs of the multi-channel data of all subjects to determine whether data was contaminated by noise and to identify broken, unresponsive channels. A semi-automatic pipeline (adjusted from the EEGLAB-based *trimOutlier* function) was used to mark and later remove the segments contaminated by artifacts from the data that corresponded to muscle activity or non-biological noise. For this purpose, we set different amplitude threshold levels for the noise detection at slow-frequency (1 – 15Hz) and high frequency (15 – 45Hz) ranges. For the slow-frequency range noise we excluded the two frontal channels (Fp1 and Fp2) for artifact detection. The individual noise threshold was defined as three standard deviations (SD) above the mean amplitude of the filtered signal in the frequency range of interest across channels, with an upper limit of 100µV. Because noise at the high-frequency range is smaller in amplitude, we set a constant amplitude threshold of 40 µV for this frequency range. Indices of bad segments were then assigned to the time series data of all scalp electrodes. Recordings for which the total bad segment length exceeded 60s were inspected visually to confirm that the marked segments were indeed contaminated by noise. An overlap between the segments detected by our algorithm and the ones marked manually by another research group using the same data [61] were compared to control for the performance of the semi-automatic algorithm. In case of a mismatch of more than 10 s, we visually inspected the recording. Subsequently, noisy segments were removed from the data. As the next step, independent component analysis (ICA, Infomax; [62]) was applied. The ICA components were visually inspected, and artifacts related to eye blinks, eye movements, heartbeat, and muscle activity were removed by excluding the corresponding components.

To exclude data periods of reduced vigilance, we only used segments of recordings that had been classified as ‘wakeful rest’ based on the Vigilance Algorithm Leipzig (VIGALL). VIGALL 2.0 [63], [64] is an automatic algorithm implemented in the Brain Vision Analzyer 2 (Brain Products GmbH, Gilching, Germany), that classifies each one second epoch of the rsEEG recording into seven categories corresponding to estimated brain arousal levels ranging from high alertness to sleep onset. The level of arousal is determined by the combination of power in different frequency bands, the EOG channel activity, and sleep spindles, as well as topographical distribution of these parameters.

### 4.3. EEG data analysis

#### 4.3.1. Aperiodic and periodic components of the power spectral density

The PSD of each channel’s data was calculated from the cleaned data using 4s Hamming windows overlapping by 50% using Welch’s method. We used the Python (version 3.6.7) implementation of the FOOOF algorithm [30] on the PSDs to estimate the slope of the 1/f decay for each channel separately: Here, broad-band PSD between 2 and 40Hz was modeled as *P*(*f*) ∼ 1/*f*^*γ*^ where *γ* is the spectral slope. FOOOF algorithm estimates the 1/f slope while excluding major peaks of the PSD (e.g. the peak in the alpha frequency range) in each channel. The same frequency range was also used in the Ouyang et al., [32] study, as opposed to the 2 - 24Hz range used in Voytek et al. [31] since this frequency range might interfere with the beta peak and result in an estimation of a steeper slope. Channels that had a good model fit (R^2^ > 0.8, which accounted for more than 99% of cases) were then used for further analysis. After subtracting the 1/f part of the spectrum from the original PSD, we performed a peak search between 7 and 13Hz to localize the alpha peak in each channel. A peak was localized if the inclination of the PSD in this frequency range exceeded 0.05 µV^2^/Hz. If the width of a peak exceeded 6Hz, it was set to 6Hz centered around the maximum of the peak. In case the peak could not be detected, we did not estimate power in that particular channel – 8.7% of participants had at least one channel in which an alpha peak could not be detected (in total 1.3% of all channels across participants had no peak in this frequency range). We then calculated alpha power as the area under the residual PSD within the frequency range between the start and end of the detected alpha peak.

We also performed a peak search between 4 and 7Hz to estimate theta peak parameters using the same criteria that were applied for the alpha peak detection. However, only ∼ 3% of participants had an oscillatory peak in this range that was consistent with findings of Caplan et al. [25] who reported the absence of an oscillatory theta peak. Because of the absence of an oscillatory peak in the theta range, we estimated theta power in a frequency range between the starting point of the alpha peak and 3Hz prior to it. Power in this range was estimated using the original PSD (containing the aperiodic component) since the subtraction of 1/f might cause negative values in the residual of the PSD. We inspected data for outliers by defining intervals of potentially ‘abnormal’ data based on the interquartile range (IQR), a method proposed by Tukey [65]. Since theta and alpha power values were positively skewed, we defined an outlier as a data value that exceeded an interval of *q*_3_ + 3 ∗ *IQR*, where *q*_3_ is the third quartile. As 1/f slope values could be both positive and negative, we have used the same interval for the positive values and an interval of *q*_1_ − 3 ∗ *IQR* for the negatively skewed values, where *q*_1_ is the first quartile. A schematic depiction of the analysis pipeline of the aperiodic and periodic components of the PSD is provided in Figure 1.

#### 4.3.2. EEG source reconstruction

To map the rsEEG data from sensor space to source space, we built individual head models for those participants that had an MRI scan (n ∼ 700) and used a standard head model for the rest of the subjects. For the individual head models, we used the T1-weighted MPRAGE images that were acquired with a 3 Tesla Verio scanner (Siemens, Erlangen, Germany) and were segmented using the Freesurfer v.5.3.0 software [66]. A 3-shell boundary element model (BEM) was constructed with Brainstorm [67], which was used to compute the leadfield matrix with OpenMEEG [68]. The standard head model was based on the ICBM152 nonlinear average head anatomy (version 2009; [69]) included within Brainstorm. Electrode positions were registered to the scalp surface of the standard head according to the 10 - 20 electrode placement standard. For individual head models, electrode positions were warped from the standard to the individual anatomy using SPM [70], after which they were projected to the individual scalp surface. In all cases the source space consisted of ca. 2000 voxels located on the cortical mantle. We constrained the orientation of the dipolar sources to be perpendicular to the cortical surface. Source reconstruction was performed using exact low-resolution brain electromagnetic tomography (eLORETA; [71]) with a regularization parameter of 0.05. The MATLAB implementation of eLORETA included in the M/EEG Toolbox of Hamburg (METH^1^) was used. Due to the small number of EEG channels (N = 31), we grouped the cortical vertices into 10 major regions that were aggregated based on the 68 regions of the Desikan-Killiany atlas [72]. We estimated parameters of the periodic and aperiodic PSD components for each vertex and plotted histograms of rsEEG parameters in each region of interest (ROI) to visually inspect them and decide on the method of vertex aggregation within an ROI. We used a geometric mean for theta and alpha power, mode for the IAF, and median for the 1/f slope. The rest of the analysis was done in the same way as described in section 2.3.1.

We compared rsEEG parameters estimated with individual as well as standard head models in 10 ROIs (see *Supplementary material Figure 1*) and since they did not differ significantly, we used a joined sample of subjects with individual (n ∼ 700) and standard (n ∼ 1000) head models for further analysis.

### 4.4. Cognition battery

#### 4.4.1. Description of cognitive tests

We used data from four cognitive tests: the Trail Making Test (TMT; [73]), Stroop test [74], Wechsler’s Memory Scale [75], and Vocabulary Knowledge Test (org. Wortschatztest, WST; [76]).

##### Trail Making Test

The TMT is a pencil-and-paper test that consists of two trails, A and B. In trail A, participants are asked to connect a series of 25 numbers in ascending order. In trail B, participants have to interchangeably connect numbers and letters in ascending and alphabetic order (i.e. 1-A-2-B…). The time it takes to complete trail A and B has been taken as a proxy for processing speed, mental flexibility, and visual-motor skills [73], [77]. We inverted the score (1/sec) for interpretability, where higher scores meant better performance.

##### Stroop Color and Word Test

The Stroop test is a neuropsychological test used to assess interference resulting from two features of a stimulus (i.e. color and meaning of the word) that are interfering with the processing of information [74]. Congruent and incongruent conditions are presented to participants. In the congruent condition, the meaning of the word displayed is consistent with the color the word is displayed in, while in the incongruent condition it is not. Reaction times in congruent and incongruent conditions as well as the accuracy measure from the correct answers from the latter condition were used in the present study. Just as in the TMT, reaction times were inverted (1/sec) for interpretability.

##### Wechsler’s Memory Scale

We used the Logical memory subscale from the WMS as a proxy for verbal episodic memory. Participants were read a short prose and scores defining episodic memory performance were aggregated based on the number of details recalled before and after a 25 minute break [75].

##### Vocabulary Knowledge Test

The WST consists of rows of words with five pseudo words that serve as distractors and a single word with semantic meaning. For each row, participants are asked to indicate the word that held meaning. The difficulty increased with consecutive rows. The score is defined as the number of correctly completed rows. The WST is used to measure crystalized intelligence.

We visually inspected cognitive scores for possible outliers and removed data in case of values associated with a typing error We then z-transformed all scores after outlier removal.

#### 4.4.2. Factor analysis of the cognition battery

Exploratory factor analysis (EFA) was used to extract latent factors underlying cognitive scales in order to reduce data dimensionality. EFA was performed with the *stats* package R (version 3.4.4). The number of latent factors was determined by the Scree plots as well as the Empirical Kaiser Criterion (EKC, [78]), according to which only the components with eigenvalues larger than one should be kept. Both, the Scree plots as well as EKC, suggested three latent factors that explained 67% of the variance in the data. These three factors were used for further statistical analyses.

### 4.5. Statistical analyses

#### 4.5.1. Relationship between resting-state EEG parameters and age

To test the relationship between the PSD components (theta power, alpha power, IAF, and 1/f slope) and their link to age, we used a mass-bivariate approach and cluster-based statistics [79] to correct for multiple comparisons across channels. For every relationship between the rsEEG parameters and age in each channel we used Pearson partial correlations. We partialed out the effects of the three other PSD variables as well as sex (measured as a bivariate choice between ‘male’ and ‘female’) and education that could have an effect on scores defining cognitive performance. Then, clusters were formed in sensor space defined as several neighboring channels (two or more) where the defined relationship passed the significance threshold of p < 0.05. If a cluster was found, the cluster t-value, estimated as a sum of t-values over electrodes that formed the cluster, was compared to a null distribution of clusters generated using the Monte Carlo method with 1000 permutations of the age values. By comparing the t-value of the original cluster with the randomly generated one, we determined the corresponding cluster p-value (*p*_*cluster*_). A cluster was considered significant if *p*_*cluster*_ ≤ 0.0125, accounting for the separate permutation tests for the four variables of interest (Bonferroni correction).

#### 4.5.2. Relationship between resting-state EEG parameters and cognition

To test the relation between the rsEEG parameters and the three cognitive factors, we used multiple linear regression (MLR) with interaction terms for age and rsEEG parameters of interest using the *lm* function implemented in R. We ran separate MLRs for each cognitive factor and each ROI due to high collinearity between brain regions. Multicollinearity might compromise the model as the effects of independent variables on the dependent one could not be reliably estimated in isolation. Therefore, each MLR model consisted of four independent variables of interest (theta power, alpha power, IAF, and 1/f slope) within a single ROI, as well as their interactions with age. Age, sex, and education were added as covariates to the models. We corrected for multiple comparisons using false discovery rate at 0.05 (FDR, [80]).

## Acknowledgements

This study has been supported by LIFE – Leipzig Research Center for Civilization Diseases, University Leipzig. LIFE is funded by means of the European Union, by the European Regional Development Fund (ERDF) and by means of the Free State of Saxony within the framework of the excellence initiative (project numbers 713-241202, 713-241202, 14505/2470, 14575/2470). Part of this work has been supported by the German Research Foundation (SCHR 774/5-1 to MLS). We thank all participants and the team at the LIFE study centre, who made this study possible.

## Data availability

Anonymized data will be made available upon request through the application procedure carried out by the LIFE-Study administration (https://life.uni-leipzig.de/de/erwachsenen_kohorten/life_adult.html).

## Code availability

Codes for data analysis are available on https://github.com/ecesnaite/periodic-aperiodic-LIFE.

## Footnotes

1 https://www.uke.de/english/departments-institutes/institutes/neurophysiology-and-pathophysiology/research/research-groups/index.html

