## Supplementary material for "Alterations in rhythmic and non-rhythmic resting-state EEG activity and their link to cognition in older age"

**Figure 1.** Comparison between resting-state EEG parameters derived at source space using standard (y axis) and individual (x axis) head models. Individual head models were reconstructed for 699 participants that had MRI scans available. EEG parameters were estimated using individual head models and compared to the parameters estimated using standard head BEM models for the same group of participants. Scatterplots show that parameters are highly comparable for theta power (A), alpha power (B), individual alpha peak frequency (C), 1/f slope of power spectral density (D) in 10 regions of interest. Abb.: l – left, r – right, Cg – cingulate cortex, OC – occipital lobe, TL – temporal lobe, PL – parietal lobe, FL – frontal lobe.

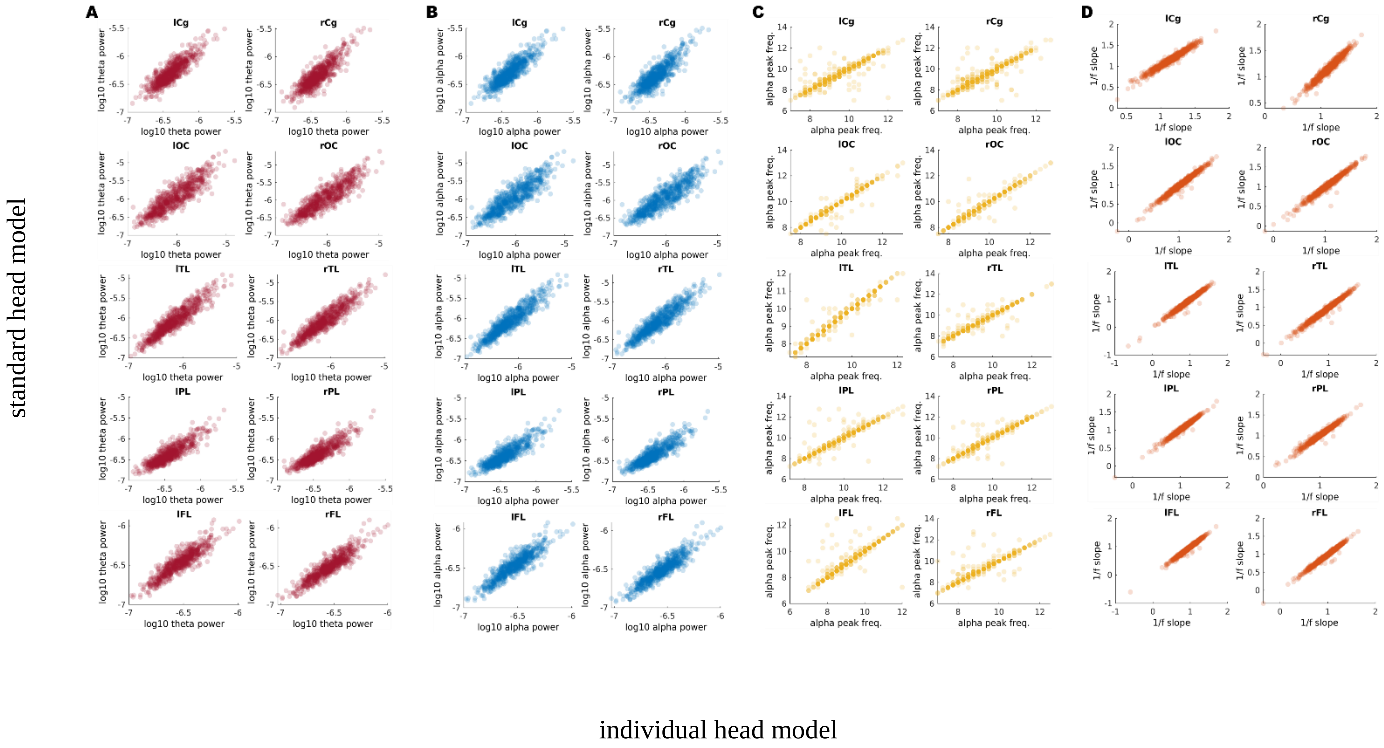

**Tables 1-10.** Output from the multiple linear models for each region of interest. The p-values are not corrected for multuple comparissons in these tables, but results that survived FDR correaction are reported in the Results section of the paper.

#### left Cignulate cortex

| | **Factor 1** | | | **Factor 2** | | | **Factor 3** | | |
|---|---|---|---|---|---|---|---|---|---|
| *Predictors* | *Estimates* | *CI* | *p* | *Estimates* | *CI* | *p* | *Estimates* | *CI* | *p* |
| Intercept | 0.06 | -0.10 – 0.23 | 0.445 | -0.36 | -0.52 – -0.21 | **<0.001** | 0.12 | -0.04 – 0.28 | 0.143 |
| IAF | 0.06 | 0.01 – 0.11 | **0.023** | -0.03 | -0.08 – 0.02 | 0.265 | 0.06 | 0.00 – 0.11 | **0.035** |
| alpha power | -0.04 | -0.10 – 0.02 | 0.187 | -0.00 | -0.06 – 0.05 | 0.961 | 0.03 | -0.02 – 0.09 | 0.237 |
| theta power | 0.00 | -0.05 – 0.06 | 0.907 | 0.00 | -0.05 – 0.06 | 0.927 | 0.02 | -0.04 – 0.07 | 0.546 |
| 1/f slope | 0.05 | -0.00 – 0.11 | 0.074 | -0.02 | -0.08 – 0.03 | 0.407 | 0.02 | -0.04 – 0.08 | 0.475 |
| age | -0.10 | -0.15 – -0.05 | **<0.001** | -0.08 | -0.13 – -0.03 | **0.001** | -0.14 | -0.19 – -0.09 | **<0.001** |
| education | 0.00 | -0.05 – 0.05 | 0.928 | 0.23 | 0.19 – 0.28 | **<0.001** | 0.08 | 0.03 – 0.13 | **0.002** |
| sex | -0.04 | -0.15 – 0.06 | 0.420 | 0.24 | 0.14 – 0.34 | **<0.001** | -0.07 | -0.17 – 0.03 | 0.180 |
| IAF:age | 0.01 | -0.04 – 0.07 | 0.585 | 0.03 | -0.02 – 0.08 | 0.269 | 0.04 | -0.01 – 0.10 | 0.098 |
| alpha power:age | -0.01 | -0.07 – 0.05 | 0.768 | 0.04 | -0.02 – 0.09 | 0.192 | -0.01 | -0.06 – 0.05 | 0.831 |
| theta power:age | -0.01 | -0.07 – 0.04 | 0.692 | 0.01 | -0.04 – 0.06 | 0.665 | 0.02 | -0.04 – 0.07 | 0.566 |
| 1/f slope:age | 0.00 | -0.05 – 0.06 | 0.863 | 0.02 | -0.03 – 0.08 | 0.378 | -0.02 | -0.07 – 0.04 | 0.541 |
| Observations | 1603 | | | 1603 | | | 1603 | | |
| $R^2$ / $R^2$ adjusted | 0.019 / 0.013 | | | 0.069 / 0.062 | | | 0.040 / 0.033 | | |

#### right Cignulate cortex

| | **Factor 1** | | | **Factor 2** | | | **Factor 3** | | |
|---|---|---|---|---|---|---|---|---|---|
| *Predictors* | *Estimates* | *CI* | *p* | *Estimates* | *CI* | *p* | *Estimates* | *CI* | *p* |
| Intercept | 0.08 | -0.09 – 0.24 | 0.366 | -0.36 | -0.51 – -0.20 | **<0.001** | 0.12 | -0.04 – 0.28 | 0.149 |
| IAF | 0.07 | 0.01 – 0.12 | **0.014** | -0.04 | -0.09 – 0.01 | 0.152 | 0.08 | 0.02 – 0.13 | **0.005** |
| alpha power | -0.01 | -0.07 – 0.05 | 0.704 | -0.02 | -0.08 – 0.04 | 0.527 | -0.00 | -0.06 – 0.06 | 0.911 |
| theta power | -0.01 | -0.07 – 0.05 | 0.775 | 0.00 | -0.05 – 0.06 | 0.940 | 0.03 | -0.03 – 0.08 | 0.382 |
| 1/f slope | 0.07 | 0.01 – 0.12 | **0.021** | -0.01 | -0.07 – 0.04 | 0.641 | 0.02 | -0.04 – 0.07 | 0.584 |
| age | -0.10 | -0.15 – -0.05 | **<0.001** | -0.08 | -0.13 – -0.03 | **0.001** | -0.15 | -0.20 – -0.10 | **<0.001** |
| education | -0.00 | -0.05 – 0.05 | 0.964 | 0.24 | 0.19 – 0.29 | **<0.001** | 0.08 | 0.03 – 0.13 | **0.003** |
| sex | -0.05 | -0.15 – 0.06 | 0.359 | 0.23 | 0.13 – 0.33 | **<0.001** | -0.08 | -0.18 – 0.03 | 0.148 |
| IAF:age | 0.02 | -0.03 – 0.07 | 0.429 | 0.03 | -0.02 – 0.08 | 0.194 | 0.03 | -0.02 – 0.08 | 0.252 |
| alpha power:age | 0.00 | -0.05 – 0.06 | 0.880 | 0.02 | -0.03 – 0.08 | 0.447 | -0.03 | -0.09 – 0.03 | 0.330 |
| theta power:age | -0.01 | -0.07 – 0.04 | 0.678 | 0.02 | -0.03 – 0.08 | 0.423 | 0.02 | -0.03 – 0.08 | 0.472 |
| 1/f slope:age | -0.01 | -0.06 – 0.05 | 0.806 | 0.01 | -0.04 – 0.06 | 0.701 | -0.02 | -0.08 – 0.03 | 0.419 |
| Observations | 1582 | | | 1582 | | | 1582 | | |
| $R^2$ / $R^2$ adjusted | 0.019 / 0.012 | | | 0.070 / 0.063 | | | 0.041 / 0.035 | | |

### left Occipital lobe

| | Factor 1 | | | Factor 2 | | | Factor 3 | | |
|---|---|---|---|---|---|---|---|---|---|
| *Predictors* | *Estimates* | *CI* | *p* | *Estimates* | *CI* | *p* | *Estimates* | *CI* | *p* |
| Intercept | 0.10 | -0.07 – 0.27 | 0.236 | -0.35 | -0.52 – -0.19 | **<0.001** | 0.10 | -0.07 – 0.27 | 0.247 |
| IAF | 0.06 | 0.00 – 0.11 | **0.035** | -0.03 | -0.08 – 0.02 | 0.276 | 0.03 | -0.02 – 0.08 | 0.292 |
| alpha power | 0.00 | -0.06 – 0.07 | 0.881 | 0.03 | -0.03 – 0.09 | 0.369 | -0.01 | -0.07 – 0.05 | 0.834 |
| theta power | 0.01 | -0.05 – 0.07 | 0.795 | -0.01 | -0.06 – 0.05 | 0.790 | 0.03 | -0.03 – 0.09 | 0.352 |
| 1/f slope | 0.00 | -0.06 – 0.06 | 0.986 | -0.03 | -0.09 – 0.03 | 0.292 | 0.01 | -0.05 – 0.07 | 0.657 |
| age | -0.10 | -0.15 – -0.05 | **<0.001** | -0.08 | -0.13 – -0.03 | **0.003** | -0.15 | -0.20 – -0.10 | **<0.001** |
| education | 0.01 | -0.04 – 0.06 | 0.716 | 0.24 | 0.19 – 0.29 | **<0.001** | 0.08 | 0.03 – 0.13 | **0.002** |
| sex | -0.07 | -0.17 – 0.04 | 0.217 | 0.23 | 0.13 – 0.34 | **<0.001** | -0.06 | -0.17 – 0.04 | 0.236 |
| IAF:age | -0.00 | -0.05 – 0.05 | 0.991 | 0.02 | -0.03 – 0.07 | 0.379 | 0.01 | -0.04 – 0.06 | 0.696 |
| alpha power:age | -0.02 | -0.08 – 0.04 | 0.540 | 0.03 | -0.03 – 0.09 | 0.315 | -0.00 | -0.06 – 0.06 | 0.950 |
| theta power:age | -0.01 | -0.06 – 0.04 | 0.720 | 0.03 | -0.02 – 0.08 | 0.275 | 0.01 | -0.04 – 0.07 | 0.613 |
| 1/f slope:age | 0.01 | -0.05 – 0.07 | 0.733 | 0.00 | -0.05 – 0.06 | 0.908 | -0.02 | -0.07 – 0.04 | 0.602 |
| Observations | 1590 | | | 1590 | | | 1590 | | |
| $R^2$ / $R^2$ adjusted | 0.018 / 0.011 | | | 0.070 / 0.064 | | | 0.035 / 0.028 | | |

### right Occipital lobe

| | Factor 1 | | | Factor 2 | | | Factor 3 | | |
|---|---|---|---|---|---|---|---|---|---|
| *Predictors* | *Estimates* | *CI* | *p* | *Estimates* | *CI* | *p* | *Estimates* | *CI* | *p* |
| Intercept | 0.09 | -0.07 – 0.26 | 0.274 | -0.36 | -0.52 – -0.20 | **<0.001** | 0.12 | -0.05 – 0.28 | 0.179 |
| IAF | 0.06 | 0.01 – 0.11 | **0.029** | -0.04 | -0.10 – 0.01 | 0.086 | 0.04 | -0.01 – 0.09 | 0.145 |
| alpha power | -0.01 | -0.07 – 0.05 | 0.695 | 0.03 | -0.03 – 0.08 | 0.398 | 0.00 | -0.06 – 0.06 | 0.927 |
| theta power | 0.00 | -0.05 – 0.06 | 0.894 | -0.01 | -0.06 – 0.05 | 0.820 | 0.02 | -0.03 – 0.08 | 0.397 |
| 1/f slope | -0.00 | -0.06 – 0.06 | 0.925 | -0.04 | -0.10 – 0.02 | 0.206 | 0.01 | -0.05 – 0.07 | 0.746 |
| age | -0.10 | -0.15 – -0.05 | **<0.001** | -0.08 | -0.13 – -0.03 | **0.002** | -0.15 | -0.20 – -0.10 | **<0.001** |
| education | 0.01 | -0.04 – 0.06 | 0.824 | 0.24 | 0.19 – 0.29 | **<0.001** | 0.08 | 0.03 – 0.13 | **0.001** |
| sex | -0.06 | -0.17 – 0.04 | 0.230 | 0.24 | 0.13 – 0.34 | **<0.001** | -0.08 | -0.18 – 0.03 | 0.159 |
| IAF:age | -0.00 | -0.05 – 0.05 | 0.911 | 0.01 | -0.04 – 0.06 | 0.640 | 0.02 | -0.04 – 0.07 | 0.540 |
| alpha power:age | -0.01 | -0.06 – 0.05 | 0.791 | 0.06 | 0.00 – 0.11 | **0.042** | 0.01 | -0.04 – 0.07 | 0.683 |
| theta power:age | -0.01 | -0.07 – 0.04 | 0.660 | 0.02 | -0.03 – 0.07 | 0.476 | 0.02 | -0.04 – 0.07 | 0.538 |
| 1/f slope:age | 0.02 | -0.04 – 0.07 | 0.552 | -0.01 | -0.06 – 0.05 | 0.764 | -0.03 | -0.09 – 0.03 | 0.286 |
| Observations | 1587 | | | 1587 | | | 1587 | | |
| $R^2$ / $R^2$ adjusted | 0.018 / 0.011 | | | 0.071 / 0.065 | | | 0.037 / 0.030 | | |

### left Temporal lobe

| | Factor 1 | | | Factor 2 | | | Factor 3 | | |
|---|---|---|---|---|---|---|---|---|---|
| *Predictors* | *Estimates* | *CI* | *p* | *Estimates* | *CI* | *p* | *Estimates* | *CI* | *p* |
| Intercept | 0.10 | -0.07 – 0.26 | 0.262 | -0.38 | -0.54 – -0.22 | **<0.001** | 0.14 | -0.03 – 0.30 | 0.112 |
| IAF | 0.07 | 0.02 – 0.12 | **0.011** | -0.04 | -0.09 – 0.01 | 0.136 | 0.08 | 0.03 – 0.14 | **0.002** |
| alpha power | -0.01 | -0.07 – 0.05 | 0.737 | 0.02 | -0.04 – 0.08 | 0.447 | -0.03 | -0.09 – 0.03 | 0.369 |
| theta power | -0.00 | -0.06 – 0.06 | 0.998 | -0.03 | -0.08 – 0.03 | 0.368 | 0.06 | -0.00 – 0.11 | 0.058 |
| 1/f slope | 0.02 | -0.04 – 0.07 | 0.558 | -0.01 | -0.07 – 0.04 | 0.639 | 0.04 | -0.02 – 0.09 | 0.201 |
| age | -0.10 | -0.15 – -0.05 | **<0.001** | -0.09 | -0.14 – -0.04 | **0.001** | -0.14 | -0.19 – -0.09 | **<0.001** |
| education | 0.00 | -0.05 – 0.05 | 0.912 | 0.24 | 0.19 – 0.29 | **<0.001** | 0.07 | 0.02 – 0.12 | **0.005** |
| sex | -0.06 | -0.17 – 0.04 | 0.242 | 0.25 | 0.15 – 0.35 | **<0.001** | -0.08 | -0.18 – 0.03 | 0.148 |
| IAF:age | -0.00 | -0.06 – 0.05 | 0.851 | 0.04 | -0.01 – 0.09 | 0.089 | 0.05 | 0.00 – 0.11 | **0.038** |
| alpha power:age | -0.04 | -0.10 – 0.02 | 0.191 | 0.03 | -0.03 – 0.08 | 0.363 | -0.02 | -0.07 – 0.04 | 0.553 |
| theta power:age | -0.01 | -0.06 – 0.05 | 0.778 | 0.02 | -0.04 – 0.07 | 0.572 | 0.02 | -0.04 – 0.07 | 0.495 |
| 1/f slope:age | 0.00 | -0.05 – 0.06 | 0.915 | 0.02 | -0.03 – 0.08 | 0.393 | 0.01 | -0.04 – 0.07 | 0.612 |
| Observations | 1591 | | | 1591 | | | 1591 | | |
| $R^2$ / $R^2$ adjusted | 0.020 / 0.013 | | | 0.074 / 0.067 | | | 0.045 / 0.038 | | |

### right Temporal lobe

| | Factor 1 | | | Factor 2 | | | Factor 3 | | |
|---|---|---|---|---|---|---|---|---|---|
| *Predictors* | *Estimates* | *CI* | *p* | *Estimates* | *CI* | *p* | *Estimates* | *CI* | *p* |
| Intercept | 0.07 | -0.10 – 0.23 | 0.444 | -0.37 | -0.53 – -0.20 | **<0.001** | 0.15 | -0.02 – 0.31 | 0.085 |
| IAF | 0.05 | 0.00 – 0.11 | **0.041** | -0.04 | -0.09 – 0.01 | 0.148 | 0.10 | 0.05 – 0.15 | **<0.001** |
| alpha power | -0.03 | -0.09 – 0.03 | 0.378 | 0.01 | -0.05 – 0.06 | 0.837 | -0.00 | -0.06 – 0.06 | 0.921 |
| theta power | 0.01 | -0.05 – 0.06 | 0.831 | -0.01 | -0.06 – 0.05 | 0.804 | 0.04 | -0.02 – 0.10 | 0.180 |
| 1/f slope | 0.02 | -0.04 – 0.07 | 0.591 | -0.02 | -0.08 – 0.03 | 0.394 | 0.04 | -0.02 – 0.10 | 0.153 |
| age | -0.11 | -0.16 – -0.06 | **<0.001** | -0.08 | -0.13 – -0.03 | **0.001** | -0.14 | -0.19 – -0.09 | **<0.001** |
| education | 0.00 | -0.05 – 0.05 | 0.899 | 0.24 | 0.19 – 0.29 | **<0.001** | 0.08 | 0.03 – 0.13 | **0.002** |
| sex | -0.04 | -0.15 – 0.06 | 0.438 | 0.24 | 0.13 – 0.34 | **<0.001** | -0.09 | -0.20 – 0.01 | 0.089 |
| IAF:age | 0.00 | -0.05 – 0.06 | 0.882 | 0.03 | -0.02 – 0.08 | 0.224 | 0.04 | -0.01 – 0.09 | 0.128 |
| alpha power:age | -0.04 | -0.10 – 0.02 | 0.195 | 0.02 | -0.04 – 0.07 | 0.523 | 0.02 | -0.04 – 0.08 | 0.450 |
| theta power:age | -0.01 | -0.06 – 0.05 | 0.739 | 0.03 | -0.03 – 0.08 | 0.308 | 0.01 | -0.05 – 0.06 | 0.841 |
| 1/f slope:age | 0.04 | -0.01 – 0.10 | 0.144 | 0.02 | -0.03 – 0.07 | 0.484 | -0.03 | -0.09 – 0.02 | 0.279 |
| Observations | 1592 | | | 1592 | | | 1592 | | |
| $R^2$ / $R^2$ adjusted | 0.020 / 0.014 | | | 0.070 / 0.064 | | | 0.046 / 0.040 | | |

### left Parietal lobe

| | Factor 1 | | | Factor 2 | | | Factor 3 | | |
|---|---|---|---|---|---|---|---|---|---|
| *Predictors* | *Estimates* | *CI* | *p* | *Estimates* | *CI* | *p* | *Estimates* | *CI* | *p* |
| Intercept | 0.06 | -0.10 – 0.23 | 0.450 | -0.38 | -0.54 – -0.21 | **<0.001** | 0.12 | -0.04 – 0.29 | 0.146 |
| IAF | 0.06 | 0.01 – 0.12 | **0.015** | -0.03 | -0.09 – 0.02 | 0.180 | 0.07 | 0.02 – 0.13 | **0.005** |
| alpha power | 0.01 | -0.05 – 0.07 | 0.803 | 0.01 | -0.05 – 0.07 | 0.791 | 0.01 | -0.05 – 0.07 | 0.779 |
| theta power | -0.02 | -0.08 – 0.04 | 0.508 | 0.01 | -0.05 – 0.07 | 0.764 | 0.02 | -0.04 – 0.08 | 0.482 |
| 1/f slope | 0.04 | -0.01 – 0.10 | 0.130 | -0.03 | -0.09 – 0.02 | 0.242 | 0.02 | -0.03 – 0.08 | 0.420 |
| age | -0.10 | -0.15 – -0.05 | **<0.001** | -0.08 | -0.13 – -0.03 | **0.002** | -0.14 | -0.19 – -0.09 | **<0.001** |
| education | 0.01 | -0.04 – 0.06 | 0.676 | 0.24 | 0.19 – 0.29 | **<0.001** | 0.08 | 0.03 – 0.13 | **0.002** |
| sex | -0.04 | -0.15 – 0.06 | 0.421 | 0.25 | 0.15 – 0.35 | **<0.001** | -0.08 | -0.18 – 0.03 | 0.155 |
| IAF:age | 0.00 | -0.05 – 0.06 | 0.890 | 0.01 | -0.04 – 0.06 | 0.733 | 0.04 | -0.01 – 0.09 | 0.141 |
| alpha power:age | -0.02 | -0.08 – 0.04 | 0.442 | 0.05 | -0.01 – 0.11 | 0.094 | -0.02 | -0.08 – 0.03 | 0.425 |
| theta power:age | -0.00 | -0.06 – 0.05 | 0.880 | 0.01 | -0.04 – 0.07 | 0.619 | 0.01 | -0.05 – 0.07 | 0.743 |
| 1/f slope:age | 0.01 | -0.05 – 0.06 | 0.795 | -0.01 | -0.06 – 0.04 | 0.665 | 0.01 | -0.04 – 0.07 | 0.666 |
| Observations | 1578 | | | 1578 | | | 1578 | | |
| $R^2$ / $R^2$ adjusted | 0.018 / 0.011 | | | 0.070 / 0.063 | | | 0.042 / 0.035 | | |

### right Parietal lobe

| | Factor 1 | | | Factor 2 | | | Factor 3 | | |
|---|---|---|---|---|---|---|---|---|---|
| *Predictors* | *Estimates* | *CI* | *p* | *Estimates* | *CI* | *p* | *Estimates* | *CI* | *p* |
| Intercept | 0.06 | -0.10 – 0.23 | 0.462 | -0.37 | -0.53 – -0.21 | **<0.001** | 0.14 | -0.03 – 0.30 | 0.103 |
| IAF | 0.06 | 0.01 – 0.11 | **0.027** | -0.03 | -0.09 – 0.02 | 0.181 | 0.07 | 0.02 – 0.12 | **0.008** |
| alpha power | -0.02 | -0.08 – 0.03 | 0.406 | 0.01 | -0.05 – 0.06 | 0.824 | 0.03 | -0.03 – 0.09 | 0.288 |
| theta power | 0.00 | -0.06 – 0.06 | 0.975 | 0.01 | -0.04 – 0.07 | 0.668 | 0.03 | -0.03 – 0.08 | 0.389 |
| 1/f slope | 0.05 | -0.00 – 0.11 | 0.064 | -0.05 | -0.10 – 0.01 | 0.081 | 0.02 | -0.04 – 0.07 | 0.586 |
| age | -0.10 | -0.15 – -0.05 | **<0.001** | -0.08 | -0.13 – -0.03 | **0.002** | -0.14 | -0.19 – -0.09 | **<0.001** |
| education | 0.00 | -0.05 – 0.05 | 0.902 | 0.24 | 0.19 – 0.29 | **<0.001** | 0.08 | 0.03 – 0.13 | **0.002** |
| sex | -0.04 | -0.15 – 0.06 | 0.413 | 0.25 | 0.15 – 0.35 | **<0.001** | -0.08 | -0.19 – 0.02 | 0.112 |
| IAF:age | -0.00 | -0.05 – 0.05 | 0.953 | 0.02 | -0.03 – 0.07 | 0.537 | 0.03 | -0.02 – 0.08 | 0.226 |
| alpha power:age | -0.03 | -0.08 – 0.03 | 0.340 | 0.06 | -0.00 – 0.11 | 0.053 | -0.02 | -0.07 – 0.04 | 0.549 |
| theta power:age | -0.01 | -0.07 – 0.05 | 0.733 | 0.02 | -0.03 – 0.07 | 0.474 | 0.02 | -0.03 – 0.08 | 0.455 |
| 1/f slope:age | 0.02 | -0.03 – 0.07 | 0.490 | 0.00 | -0.05 – 0.06 | 0.929 | -0.05 | -0.10 – 0.00 | 0.074 |
| Observations | 1586 | | | 1586 | | | 1586 | | |
| $R^2$ / $R^2$ adjusted | 0.019 / 0.012 | | | 0.072 / 0.066 | | | 0.044 / 0.037 | | |

### left Frontal lobe

| | Factor 1 | | | Factor 2 | | | Factor 3 | | |
|---|---|---|---|---|---|---|---|---|---|
| *Predictors* | *Estimates* | *CI* | *p* | *Estimates* | *CI* | *p* | *Estimates* | *CI* | *p* |
| Intercept | 0.02 | -0.15 – 0.19 | 0.781 | -0.35 | -0.52 – -0.19 | **<0.001** | 0.10 | -0.07 – 0.27 | 0.256 |
| IAF | 0.06 | 0.01 – 0.11 | **0.028** | -0.01 | -0.06 – 0.04 | 0.621 | 0.06 | 0.00 – 0.11 | **0.033** |
| alpha power | -0.06 | -0.12 – -0.00 | **0.037** | 0.01 | -0.05 – 0.07 | 0.767 | 0.01 | -0.05 – 0.07 | 0.847 |
| theta power | 0.02 | -0.03 – 0.08 | 0.418 | -0.01 | -0.07 – 0.04 | 0.623 | 0.03 | -0.02 – 0.09 | 0.265 |
| 1/f slope | 0.04 | -0.02 – 0.09 | 0.167 | -0.02 | -0.07 – 0.04 | 0.540 | 0.04 | -0.01 – 0.09 | 0.153 |
| age | -0.10 | -0.15 – -0.05 | **<0.001** | -0.08 | -0.12 – -0.03 | **0.003** | -0.14 | -0.19 – -0.09 | **<0.001** |
| education | 0.01 | -0.04 – 0.06 | 0.732 | 0.24 | 0.19 – 0.29 | **<0.001** | 0.08 | 0.03 – 0.13 | **0.003** |
| sex | -0.02 | -0.13 – 0.09 | 0.738 | 0.23 | 0.13 – 0.34 | **<0.001** | -0.06 | -0.17 – 0.05 | 0.270 |
| IAF:age | 0.00 | -0.05 – 0.05 | 0.875 | 0.03 | -0.02 – 0.08 | 0.219 | 0.01 | -0.04 – 0.06 | 0.818 |
| alpha power:age | -0.02 | -0.08 – 0.04 | 0.454 | 0.04 | -0.02 – 0.09 | 0.232 | -0.01 | -0.07 – 0.05 | 0.715 |
| theta power:age | -0.00 | -0.05 – 0.05 | 0.993 | 0.02 | -0.04 – 0.07 | 0.515 | 0.01 | -0.04 – 0.07 | 0.623 |
| 1/f slope:age | -0.02 | -0.07 – 0.03 | 0.440 | 0.04 | -0.02 – 0.09 | 0.174 | 0.00 | -0.05 – 0.05 | 0.982 |
| Observations | 1595 | | | 1595 | | | 1595 | | |
| $R^2$ / $R^2$ adjusted | 0.022 / 0.015 | | | 0.069 / 0.062 | | | 0.037 / 0.030 | | |

### right Frontal lobe

| | Factor 1 | | | Factor 2 | | | Factor 3 | | |
|---|---|---|---|---|---|---|---|---|---|
| *Predictors* | *Estimates* | *CI* | *p* | *Estimates* | *CI* | *p* | *Estimates* | *CI* | *p* |
| Intercept | 0.01 | -0.16 – 0.18 | 0.880 | -0.35 | -0.51 – -0.18 | **<0.001** | 0.12 | -0.05 – 0.29 | 0.164 |
| IAF | 0.05 | -0.00 – 0.10 | 0.060 | -0.04 | -0.09 – 0.01 | 0.166 | 0.07 | 0.02 – 0.12 | **0.006** |
| alpha power | -0.08 | -0.14 – -0.02 | **0.010** | 0.03 | -0.03 – 0.09 | 0.317 | 0.02 | -0.04 – 0.08 | 0.476 |
| theta power | 0.02 | -0.04 – 0.08 | 0.483 | -0.01 | -0.07 – 0.04 | 0.614 | 0.03 | -0.03 – 0.08 | 0.328 |
| 1/f slope | 0.07 | 0.01 – 0.12 | **0.013** | -0.02 | -0.07 – 0.03 | 0.433 | 0.03 | -0.03 – 0.08 | 0.335 |
| age | -0.10 | -0.16 – -0.05 | **<0.001** | -0.08 | -0.13 – -0.03 | **0.002** | -0.14 | -0.19 – -0.09 | **<0.001** |
| education | 0.01 | -0.04 – 0.06 | 0.770 | 0.24 | 0.19 – 0.29 | **<0.001** | 0.08 | 0.03 – 0.13 | **0.002** |
| sex | -0.01 | -0.12 – 0.10 | 0.857 | 0.23 | 0.12 – 0.33 | **<0.001** | -0.07 | -0.18 – 0.03 | 0.185 |
| IAF:age | 0.02 | -0.03 – 0.07 | 0.441 | 0.03 | -0.02 – 0.08 | 0.271 | 0.02 | -0.03 – 0.07 | 0.521 |
| alpha power:age | -0.01 | -0.07 – 0.05 | 0.705 | 0.03 | -0.03 – 0.08 | 0.370 | -0.00 | -0.06 – 0.06 | 0.900 |
| theta power:age | -0.00 | -0.06 – 0.05 | 0.914 | 0.02 | -0.03 – 0.07 | 0.497 | 0.01 | -0.04 – 0.07 | 0.693 |
| 1/f slope:age | -0.02 | -0.07 – 0.03 | 0.494 | 0.02 | -0.03 – 0.07 | 0.395 | -0.02 | -0.07 – 0.03 | 0.474 |
| Observations | 1592 | | | 1592 | | | 1592 | | |
| $R^2$ / $R^2$ adjusted | 0.025 / 0.018 | | | 0.070 / 0.063 | | | 0.039 / 0.032 | | |
